# Predator odor (TMT) exposure potentiates interoceptive sensitivity to alcohol and increases GABAergic gene expression in the anterior insular cortex and nucleus accumbens in male rats

**DOI:** 10.1101/2022.02.16.480725

**Authors:** Ryan E. Tyler, Maya N. Bluitt, Kalynn Van Voorhies, Laura C. Ornelas, Benjamin Z.S. Weinberg, Joyce Besheer

## Abstract

Post-traumatic stress disorder (PTSD) confers enhanced vulnerability to develop comorbid alcohol use disorder (AUD). Exposure to the scent of a predator, such as the fox odor TMT, has been used to model a traumatic stressor with relevance to PTSD symptomatology. Alcohol produces distinct interoceptive (subjective) effects that may influence vulnerability to problem drinking and AUD. As such, understanding the lasting impact of stressor on sensitivity to the interoceptive effects of alcohol is clinically relevant. The present study used a 2-lever, operant drug discrimination procedure to train male, Long-Evans rats to discriminate the interoceptive effects of alcohol (2 g/kg, i.g.) from water. Upon stable performance, rats underwent a 15-min exposure to TMT. Two weeks later, an alcohol dose-response curve was conducted to evaluate the lasting effects of the TMT stressor on the interoceptive effects of alcohol. The TMT group showed a leftward shift in ED_50_ of the dose response curve compared to controls, reflecting potentiated interoceptive sensitivity to alcohol. TMT exposure did not affect response rate. GABAergic signaling in both the anterior insular cortex (aIC) and the nucleus accumbens (Acb) is involved in the interoceptive effects of alcohol and stressor-induced adaptations. As such, follow-up experiments in alcohol-naïve rats examined neuronal activation (as measured by c-Fos immunoreactivity) following TMT and showed that TMT exposure increased c-Fos expression in the aIC and the nucleus accumbens core (AcbC). 2 weeks after TMT exposure, *Gad-1* gene expression was elevated in the aIC and *Gat-1* was increased in the Acb compared to controls. Lastly, the alcohol discrimination and alcohol-naïve groups displayed dramatic differences in stress reactive behaviors during the TMT exposure, suggesting that alcohol exposure may alter the behavioral response to predator odor. Together, these data suggest that predator odor stressor results in potentiated sensitivity to alcohol possibly through GABAergic adaptations in the aIC and Acb, which may be relevant to understanding PTSD-AUD comorbidity.

## Introduction

Post-traumatic stress disorder (PTSD) is a neuropsychiatric disorder that develops in some individuals after experiencing or witnessing a traumatic stress event [1]. PTSD is defined by four symptom clusters, including re-experiencing (flashbacks, unwanted memories), avoidance (external reminders), hyperarousal/hypervigilance, and negative mood/thoughts [2]. Additionally, PTSD enhances one’s risk of developing a co-morbid alcohol use disorder (AUD) [3]. Unfortunately, PTSD and AUD are lasting and debilitating disorders for which only limited treatment options exist [1]. Therefore, the goal of the present work was to begin to study how stressor events impact sensitivity to the effects of alcohol, as this may be a factor in PTSD-AUD comorbidity.

The interoceptive effects of alcohol refer to the internal bodily state induced by alcohol consumption. Because interoception (e.g., process of sensing, integrating bodily signals) plays an important role in guiding behavior, the interoceptive effects produced by alcohol can influence alcohol intake in both alcohol-dependent and non-dependent individuals by signaling satiety or influencing more drinking [4–11]. Operant drug discrimination is a well-characterized tool for investigating the interoceptive stimulus effect of drugs in preclinical models [12–16]. Using these procedures, the interoceptive drug cue is trained as a discriminative stimulus that guides a specific behavior (e.g., lever press). For example, in the 2-lever operant paradigm used here, responses on one lever (e.g., left lever) are reinforced with sucrose on drug sessions (e.g., following alcohol administration), and responses on the other lever (e.g., right lever) are reinforced with sucrose on vehicle sessions (e.g., following water administration). After many training sessions, the interoceptive state (alcohol stimulus or no alcohol stimulus) informs the correct lever selection. The percent of alcohol-appropriate lever selection serves as a quantitative index of interoceptive sensitivity to alcohol. As such, this methodology is useful for the investigation of neuropharmacological and circuitry mechanisms that underlie interoceptive states [12, 17]. Moreover, such studies have demonstrated that GABA_A_ and NMDA receptors, in part, modulate the interoceptive effects of alcohol [6, 14–16, 18–21].

Exposure to predator odors (e.g., bobcat urine, fox feces, soiled litter) in rodents have been employed to model aspects of traumatic stress exposure with relevance to some aspects of PTSD [22–40]. The predator odor 2,5-dihydro-2,4,5-trimethylthiazoline (TMT) is a synthetic component of fox feces that has been shown to produce freezing, avoidance and defensive digging behaviors as well as molecular and physiological adaptations in the brain [34, 35, 39–42]. Additionally, several studies have reported increased alcohol drinking following predator odor exposure [40, 43–46], which may be influenced by a change to the interoceptive effects of alcohol. Indeed, prior work has shown that 7-day exposure to the stress-related hormone corticosterone blunts sensitivity to the interoceptive effects of alcohol [47–49]. However, to date the impact of predator odor exposure on alcohol interoceptive effects has yet to be investigated.

The primary goal of the present work was to determine whether TMT exposure affects interoceptive sensitivity to alcohol in male rats trained to discriminate the interoceptive effects of alcohol (2 g/kg) vs. water. Clinical studies have found a strong link between GABAergic gene expression changes and PTSD in post-mortem brain tissue [50] and alcohol interoceptive effects are in part mediated by GABA receptor activation by alcohol [7, 14, 15, 17, 51, 52].

Furthermore, the anterior insular cortex (aIC) and the nucleus accumbens (Acb) have both been identified as key brain regions in mediating the interoceptive stimulus effects of alcohol [53–57]. Therefore, in naïve male rats, c-Fos immunoreactivity and GABAergic gene expression analyses were conducted in the anterior insular cortex (aIC) and the nucleus accumbens (Acb). c-Fos was measured immediately following TMT exposure (100 min), and gene expression was evaluated 2 weeks after the exposure, which is the same time point interoceptive sensitivity to alcohol was evaluated. Finally, another feature of this study was that by using alcohol-naïve rats for the gene expression analyses, we were able to investigate whether alcohol experience/history influences stress reactive behaviors during the TMT exposure.

## Methods

### Animals

Male, Long-Evans rats (N=43; Envigo, Indianapolis, IN) were used for all experiments. Rats arrived at 7 weeks and were housed individually. The vivarium was maintained on a 12-hr light/dark cycle and all experiments were conducted during the light cycle. All rats were handled for at least 1 minute daily for 1 week prior to beginning experiments. For Experiment 1 (the drug discrimination experiment), food intake was restricted to maintain body weight (325-340 g). For Experiment 2 (c-Fos experiment) and Experiment 3 (gene expression experiment), rats had ad libitum access to food and were alcohol naive. All rats had ad libitum access to water. Rats were under continuous care and monitoring by veterinary staff from the Division of Comparative Medicine at UNC-Chapel Hill. All procedures were conducted in accordance with the NIH Guide to Care and Use of Laboratory Animals and institutional guidelines.

### Experiments (detailed methodology included after the experiment descriptions)

#### Experiment 1 – The effect of TMT exposure on alcohol interoceptive sensitivity

Experiment 1 started with daily training of rats to discriminate the interoceptive stimulus effects of alcohol (2 g/kg, i.g.) from water. Upon completion of training criteria (see Discrimination Training; approximately 10 months), a single water and a single alcohol (2 g/kg) test session were conducted on consecutive days (see Discrimination Testing) to confirm stimulus control by alcohol. The following day, rats underwent the 15-min TMT exposure (N=7 CTRL v. N=6 TMT). Rats were then returned to the home cage for 2 weeks. Next, 4 different doses of alcohol (0.0, 0.5, 1.0, 2.0 g/kg, i.g.) were administered prior to a test session on 4 consecutive days to obtain a dose-response curve. Doses were counterbalanced by day and all rats received all alcohol doses.

#### Experiment 2 – The effect of TMT exposure on c-Fos immunoreactivity

Experiment 2 is the c-Fos experiment in which alcohol-naïve rats were exposed to TMT for 15 min (N=7 CTRL v. N=8 TMT). Experiment 2 did not include bedding in the test chamber and was not video recorded. 100 min after the end of the exposure, rats were sacrificed, and brains collected for quantification of c-Fos immunoreactivity (IHC) in the anterior insular cortex (aIC) and the nucleus accumbens core (AcbC).

#### Experiment 3 – The effects of TMT exposure on GABAergic gene expression

Experiment 3 is the gene expression experiment in alcohol-naïve rats. Identical to Exp. 1, rats were exposed to TMT for 15-min and returned to their home cage for 2 weeks (N=4 CTRL v. N=11 TMT). A larger sample size was used for the TMT group because prior data showed that TMT exposure results in distinct sub-groups within the TMT-exposed group that differ in brain gene expression [34]. After the 2 weeks, rats were sacrificed, and the aIC and Acb (targeting the core) brain regions were punched and collected for qRT-PCR analyses of *Gad-1, Gad-2*, and *Gat-1* expression. See Table 2 for gene expression details.

### Methodological details

#### Operant Drug Discrimination

##### Apparatus

The chambers (Med Associates, Georgia, VT; measuring 31 × 32 × 24 cm) in which all drug discrimination training and testing occurred have been previously detailed [54]. Briefly, on the right-side wall of the chambers, two retractable levers were on either side of a liquid dipper receptacle. A stimulus light was located above each dipper. Completion of an FR10 lever press/response activated the dipper to present 0.1 mL of sucrose (10% w/v) for 4 seconds. Chambers were inside of a sound-attenuating cubicle with an exhaust fan. Med Associates program was used to control sessions and record data.

##### Discrimination Training

Training sessions occurred daily (Monday-Friday) and are similar to our previously published work [51, 54, 58, 59]. Rats were administered water or alcohol (2 g/kg, i.g.), and then placed in the operant chambers for a 20 min time out period. After this period, both levers (right and left) were introduced to the chamber and the stimulus light was illuminated to indicate the start of the 15-min training session. During an alcohol session, completion of a fixed ratio 10 (FR10) on the alcohol-appropriate lever resulted in delivery of sucrose reinforcer (10% w/v, 0.1 mL). Alternatively, during water sessions, completion of FR10 on the water-appropriate lever resulted in delivery of the sucrose reinforcer. During both training sessions, responding on the inappropriate lever was recorded but did not produce a programmed consequence. The alcohol and water appropriate levers were counterbalanced across rats. Training days varied on a double alternation schedule (alcohol, alcohol, water, water,…). Testing began when criteria were met such that the percentage of appropriate lever responses before the first reinforcer and the average of the entire session was >80% for at least 8 out of 10 consecutive days.

##### Discrimination Testing

Test sessions were similar to training sessions except they were 2-min in duration (after the 20 min delay) and FR10 on either lever resulted in sucrose delivery so as not to bias lever selection and to enable response rate quantification (measure of nonspecific motor effects). Identical to training sessions, a 20-min delay period elapsed before the start of the 2-min test session.

#### Predator Odor Exposure

##### TMT Exposure

2,5-dihydro-2,4,5-trimethylthiazoline (TMT) exposure test chambers and experimental set-up were identical to those used in [39, 40, 60]. Briefly, rats were transported from the vivarium in the home cage to a separate, well-ventilated room that contained the test chambers in which rats were exposed to TMT (45.72 × 17.78 × 21.59 cm; UNC Instrument Shop, Chapel Hill, NC). Only one rat was placed in each chamber. The length of the back wall of the test chambers was opaque white with two opaque black side walls and a clear, plexiglass front wall to enable video recordings and a clear sliding lid. The bottom of the chamber was covered with a layer of white bedding (approximately 1200 ml) [39, 40, 60] except for the c-Fos experiment (Experiment 2) in which no bedding was present. A small, metal basket was hung on the right-side wall (17.8 cm above the floor) to hold a piece of filter paper. 10 μL of TMT (2,5-dihydro-2,4,5-trimethylthiazoline) - or water for controls - was pipetted onto the filter paper in the metal basket immediately prior to putting the rat in the chamber. The control group was always run before the TMT group to prevent odor contamination. The odor exposure session lasted 15 mins and was video recorded (except for Experiment 2) for evaluation of behavior using ANY-maze Video Tracking System (Version 6.12, Stoelting Co. Wood Dale, IL).

#### Brain tissue collection and sectioning

For the c-Fos experiment (Experiment 2), 100 min after the end of the TMT exposure, rats were injected with pentobarbital (100 mg/kg, i.p) prior to perfusion with 0.1 M PBS (4 °C, pH = 7.4), and then followed by 4% paraformaldehyde (PFA; 4 °C, pH = 7.4) fixation. Brains were removed and stored in 4% PFA for 24 h at 4 °C. Next, brains were rinsed with 0.1 M PBS before being moved to a 30% sucrose in 0.1 M PBS solution. 80-μm coronal sections were collected using a freezing microtome. Table 1 shows regions of quantification and representative images. Sections were stored in cryoprotectant at −20 °C until starting immunohistochemistry (IHC).

**Table 1.**
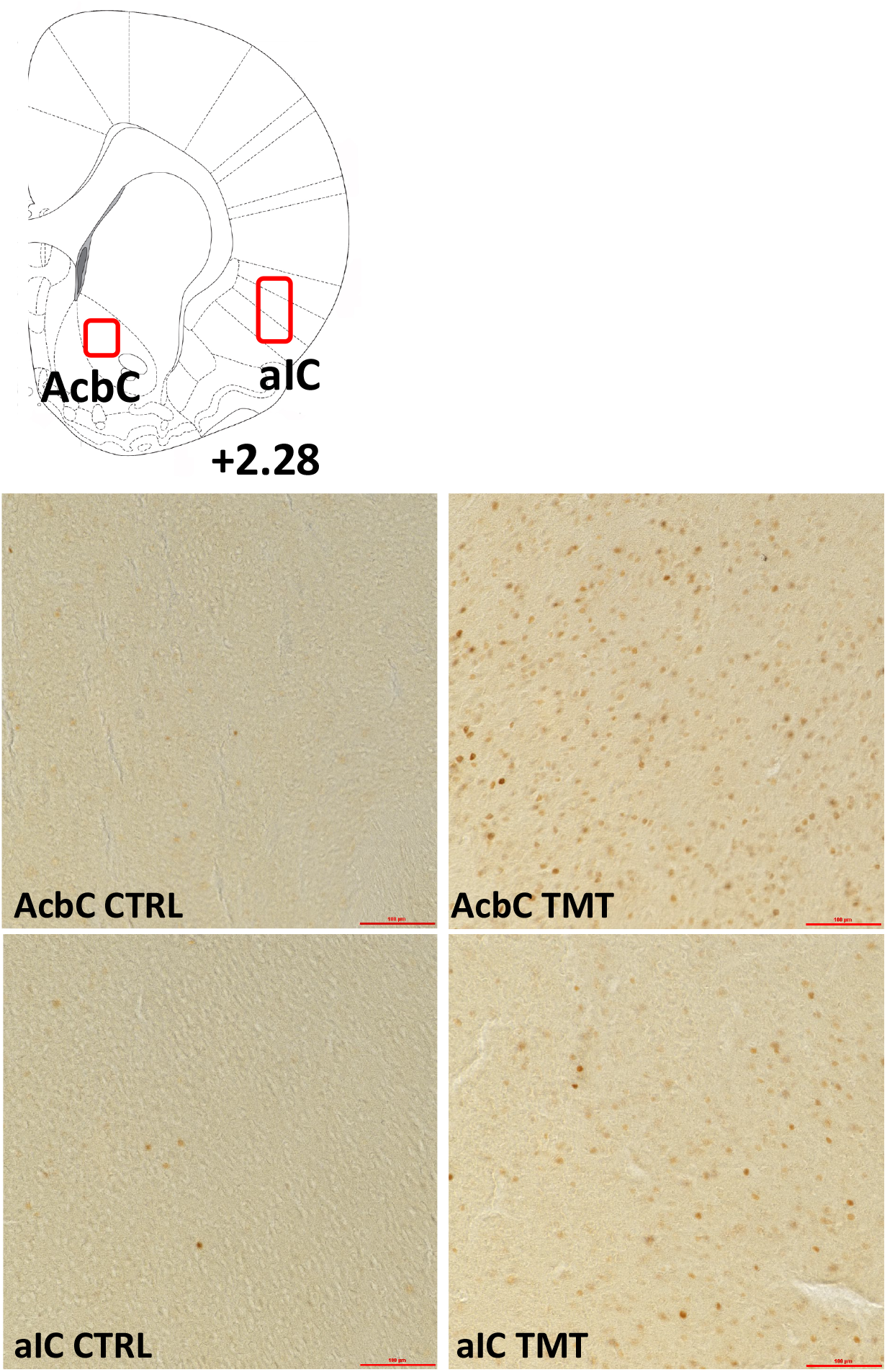
c-Fos Images (Experiment 2)

For the PCR experiment (Experiment 3), two weeks after the TMT exposure, rats were sacrificed for qPCR experiments. Rats were anesthetized in 5% isoflurane immediately before brain extraction. Brains were rapidly extracted, and flash frozen with isopentane (Sigma-Aldrich, MI). Brain tissue was stored at −80 °C until brain region sectioning. Brains were sectioned on a cryostat (−20 °C) until reaching a predetermined bregma for each region of interest (ROI) according to [61]. Next, a micropunch tool was used to extract tissue specific to the brain region of interest. These are illustrated in Table 2. Brain tissue punches were then stored at −80°C until RNA extraction.

**Table 2.**
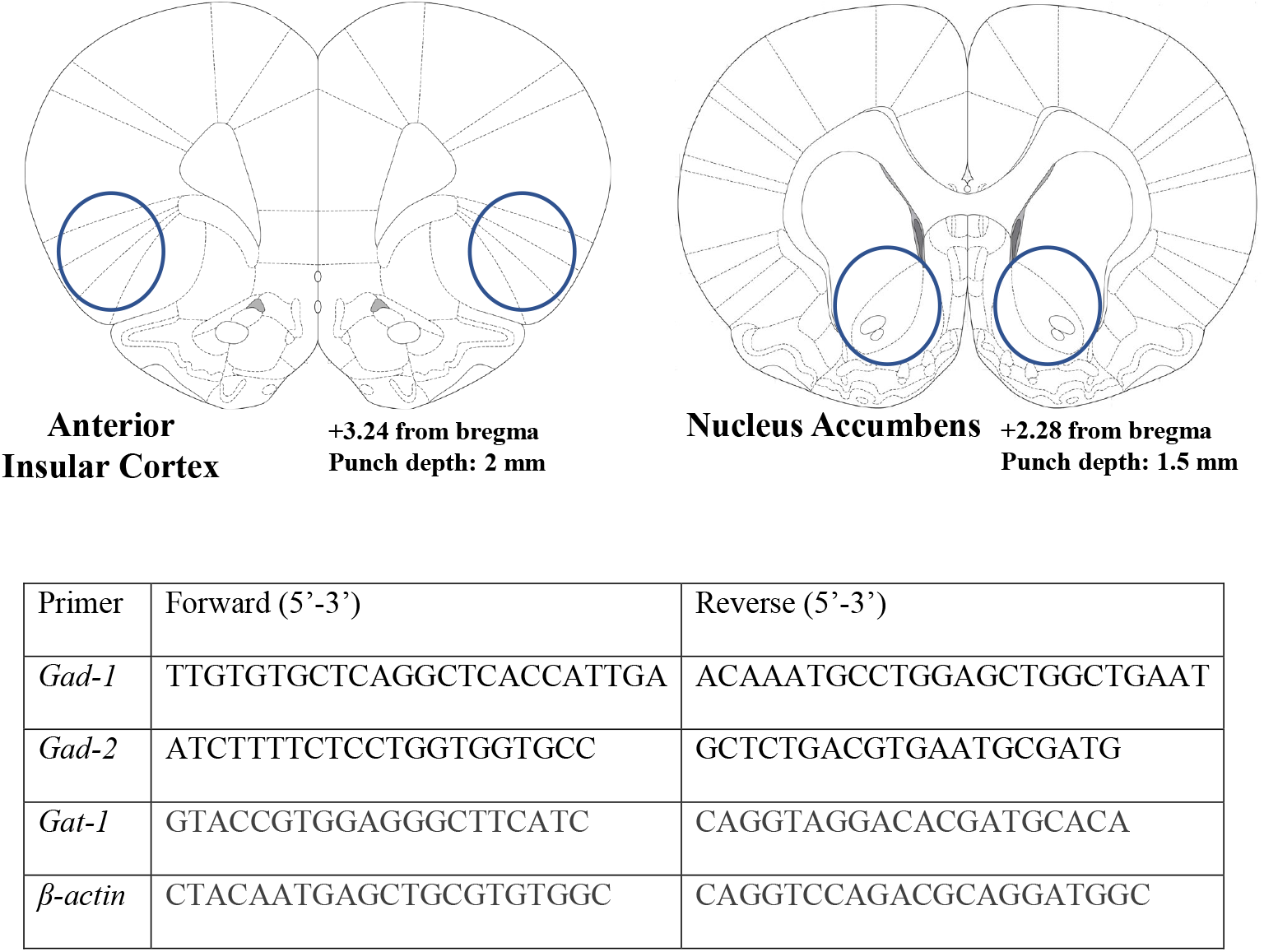
Gene expression experiment (Experiment 3)

#### c-Fos Immunohistochemistry and quantification

Coronal sections (free floating) were rinsed in PBS (0.1 M) before a 5-min wash in 1% hydrogen peroxide. Next, sections were blocked in 3% normal goat serum (NGS; Vector Labs, Burlingame, CA) in 0.3% Triton X-100 for 2 h. Sections were then incubated in rabbit anti-c-Fos antibody (1:4000 in 3% NGS + 0.1% Triton; Synaptic Systems, Gottingen, Germany; Lot# 226003/3-45) for 16 h at 4 °C. This was followed by an incubation with biotinylated goat anti-rabbit secondary antibody (1:200 in 3% NGS + 0.1% Triton X-100; Vector Labs, Burlingame, CA; Lot# 140893). Next, sections were incubated in Vectastain Elite ABC HRP (Vector labs, Burlingame, CA). Lastly, brain sections were treated with diaminobenzidine (Sigma-Aldrich, St. Louis, MO) and then mounted on glass slides.

Brain section images were obtained using the Olympus CX41 light microscope (Olympus America, Center Valley, PA). The analyses were conducted with Image-Pro Premier image analysis software (Media Cybernetics, Rockville, MD). Bilateral immunoreactivity data (c-Fos-positive cells/mm^2^; [43, 54]) were taken from a minimum of 2 sections/brain region/animal (e.g., 4 data points per animal), conducted by an experimenter blind to animal groups. These data were then averaged to obtain one value for each subject as in [43, 48, 54]. The regions examined were the anterior insular cortex (aIC, bregma +0.84 to +2.28) and the nucleus accumbens core (AcbC, bregma +1.62 to +2.52). Table 1 shows regions of quantification and representative images at 20x magnification.

#### Gene Expression Analyses

##### RNA Extraction

The RNeasy Mini Kit (Qiagen, Venlo, Netherlands) was used to extract RNA from brain tissue according to the manufacturer’s instructions. A Spectrophotometer (Nanodrop 2000, ThermoScientific) was used to determine RNA concentration and purity for each sample.

##### Reverse Transcription

The SuperScript™ III First-Strand Synthesis System (ThermoFisher Scientific) was used to reverse transcribed RNA into cDNA according to the manufacturer’s instructions. Next, all samples were diluted 1:5 with water and stored at −20°C before RT-PCR experiments.

##### RT-PCR

All experiments used the QuantStudio3 PCR machine (ThermoFisher). All samples were run in triplicate with a 96-well plate, using 10 μL total volume per well with the following components: PowerUp Syber green dye (ThermoFisher, containing ROX dye for passive reference), forward and reverse primers (Eton Biosciences Inc., NC) and cDNA template. The PCR was run with an initial activation for 10 minutes at 95 °C, followed by 40 cycles of the following: denaturation (95°C for 15 seconds), annealing/extension (60°C for 60 seconds). Melt curves were run for all genes to verify synthesis of only one amplicon. *β-actin* served as the housekeeper gene. Table 2 displays all primer sequences.

## Data Analysis

### Operant Alcohol Discrimination

Response accuracy was evaluated as the percentage of alcohol-appropriate lever response upon delivery of the first sucrose reinforcer (i.e. the first FR10; prior to feedback from sucrose delivery). Full expression of the discriminative effects of alcohol was defined as ≥80% alcohol-appropriate responses. No substitution was defined as <40% alcohol-appropriate responding. Partial substitution was defined as between 40% and 80% alcohol-appropriate responding [54, 62, 63]. The response rate was calculated as total lever responses per min and served as an index of motor activity. The ED_50_ was calculated for each rat for the dose-response curve of alcohol-appropriate lever responses (%) as the dose of alcohol which produced 50% alcohol-appropriate lever responding. The 2 doses of alcohol which encompassed 50% alcohol-appropriate responding were used to create a simple linear regression. Next, the x-value (alcohol dose) when y was set to 50% was calculated as the ED_50_. For all dose-response curves, a two-way ANOVA was used with dose as a within-subject factor and TMT exposure as a between subject factor. For ED_50_, a one-tailed, unpaired student’s t-test was used to compare the control to the TMT group. A one-tailed test was used because the direction (increase) of the effect from TMT exposure was already known from the two-way ANOVA. P ≤ 0.05 was set for significance.

### c-Fos quantification

The average number of c-Fos positive cells were compared between the TMT group and the control group using a two-tailed, unpaired student’s t-test. P ≤ 0.05 was set for significance.

### Gene Expression

The ΔΔCt method was used to calculate fold change relative to controls [64]. Fold changes were normalized such that the average control fold change was equal to 1. A two-tailed, unpaired student’s t-test with Welch’s correction was used to compare the TMT to the control group. A Welch’s correction was used due to the difference in sample size between the control and TMT groups (N=4 CTRL v. 11 TMT). In the nucleus accumbens (Acb), 2 samples were lost due to low RNA concentration, resulting in N=4 v. N=9 for the Acb. P ≤ 0.05 was set for significance.

### TMT Exposure Behavior

Analysis of TMT exposure behaviors are identical to [39, 40, 60]. The length of the TMT exposure chamber (rectangular) was divided in the middle into two compartments for analysis (TMT side and non-TMT side) using ANY-maze software. A metal basket containing TMT (10μL on a piece of filter paper) was located on the far end of one side of the chamber - termed “the TMT side.” The dependent measures evaluated included time spent digging (sec), time spent immobile (sec), time spent grooming (sec), distance traveled (meters), time spent on the TMT side of the chamber (sec), and midline crossings (the number of times the animal crossed between the TMT and non-TMT side). Immobility was operationally defined as the absence of movement except respiration for at least 2 seconds or more and determined automatically using ANYmaze software (see [39]). Time spent digging in the bedding and time spent grooming (see [39, 60]) were quantified manually by an experimenter blind to experimental conditions. For all behaviors, a two-tailed, unpaired student’s t-test was used to compare the control and TMT group. A Welch’s correction was used for the gene expression experiment (Experiment 3) due to the difference in sample size between groups. P ≤ 0.05 was set for significance.

## Results

### Timeline of Experiments

Figure 1 shows the experimental timeline for all three experiments: The operant drug discrimination (Experiment 1, Fig. 1A), c-Fos (Experiment 2, Fig. 1B), and gene expression (Experiment 3, Fig. 1C) experiments.

**Figure 1.**
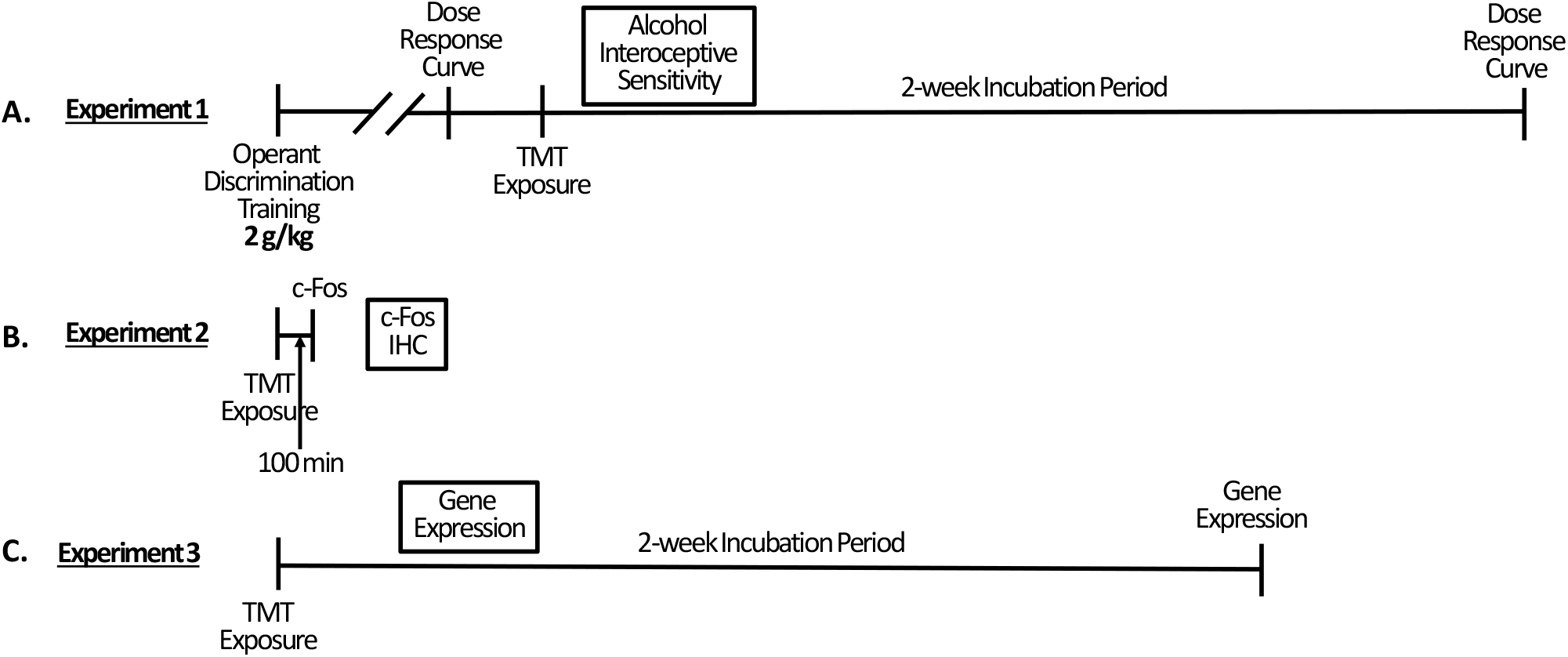
Experimental Timelines. (A) Operant drug discrimination experiment (Experiment 1). (B) c-Fos experiment (Experiment 2). (C) Gene expression experiment (Experiment 3).

#### Operant lever response is under the control of the alcohol training dose (2 g/kg)

Prior to TMT exposure (Figure 2), rats underwent 2 test sessions, one water test and one alcohol (training dose, 2 g/kg) test to confirm high accuracy performance. For the percent alcohol-appropriate lever response (Fig. 2A), there was a main effect of alcohol dose (F (1, 11) = 1475, p<0.0001) and no effect of group. Rats showed full substitution for the alcohol training dose (2 g/kg) and no substitution at 0 g/kg. For response rate (Fig. 2B), there was a main effect of alcohol dose (F (1, 11) = 14.95, p=0.002) and no effect of group. Together, these data confirm appropriate discriminative stimulus control in both groups, and no difference in accuracy performance between groups prior to TMT exposure.

**Figure 2.**
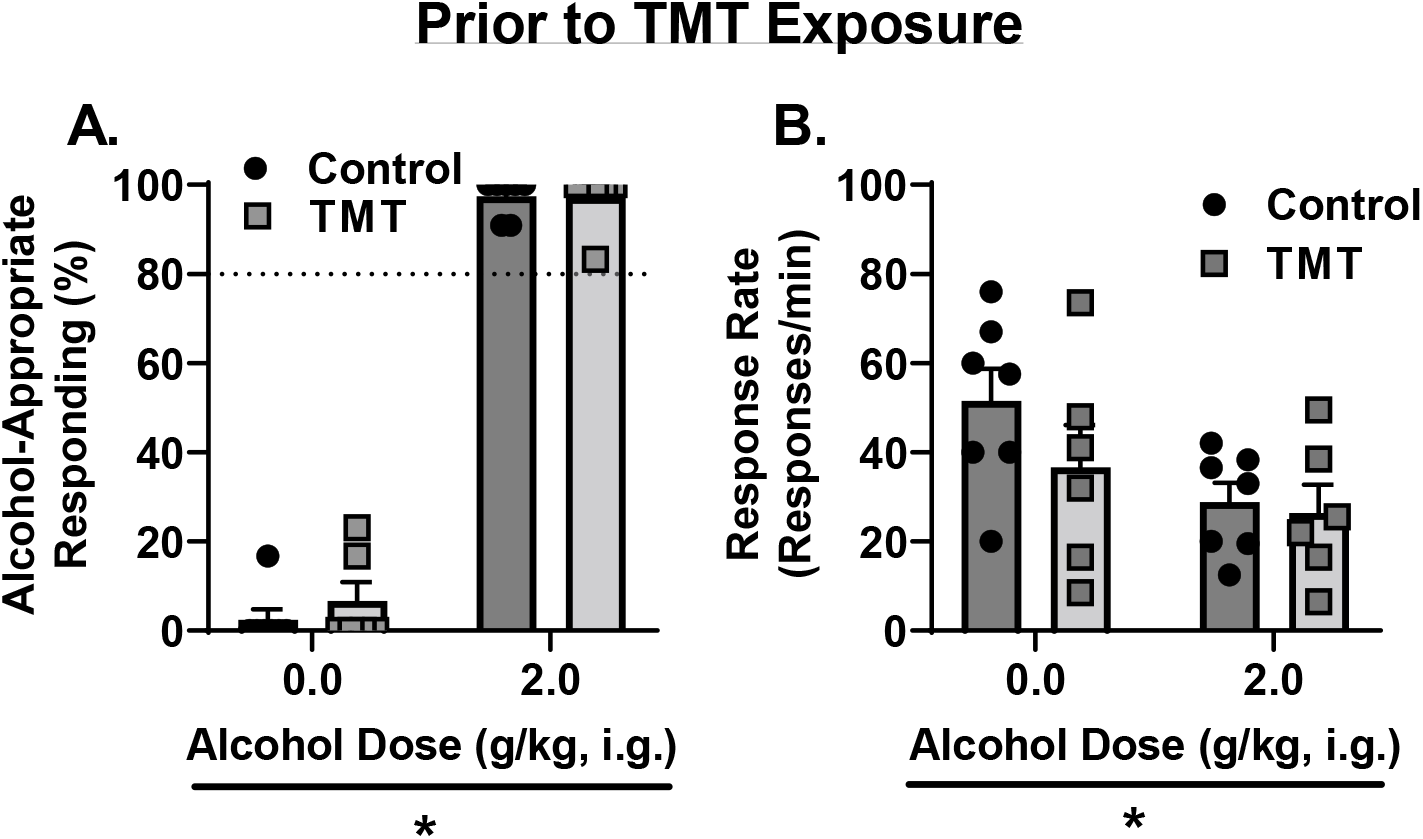
Potentiated sensitivity to the interoceptive effects of alcohol 2 weeks after TMT exposure. Prior to TMT exposure, (A) rats showed appropriate stimulus control with high alcohol-appropriate lever responses following alcohol administration and low alcohol-appropriate lever responses following water. (B) Response rate was lower following alcohol than water administration. Dotted lines at 80% indicate full substitution for the 2 g/kg alcohol training dose. p<0.05

#### TMT exposure potentiates interoceptive sensitivity to alcohol 2 weeks after stressor exposure

Two weeks after TMT exposure, for the percent alcohol-appropriate lever responses (Fig. 3A), there was a main effect of TMT exposure (F (1, 11) = 5.52, p=0.038) and alcohol dose (F (3, 33) = 47.18, p<0.0001), and a trend towards an interaction effect (F (3, 33) = 2.41, p=0.08). The TMT group showed higher alcohol-appropriate responses than the Control group. At 1 g/kg alcohol, the TMT group displayed partial substitution for the training dose (2 g/kg), but the Control group showed no substitution for the training dose. At the training dose, both TMT and Control groups showed full substitution. Analysis of the ED_50_ on the dose-response curve for alcohol appropriate responses (%) showed a decrease in the TMT group compared to controls (Fig. 3B, t(11)=1.78, p=0.05), reflecting a leftward shift of the dose-response curve. For response rate (Fig. 3C), there was a main effect of alcohol dose (F (3, 33) = 5.04, p=0.005), but no effect of TMT exposure or interaction effect. Together, these data indicate that 2 weeks following TMT exposure, rats displayed potentiated interoceptive sensitivity to alcohol, and this potentiation was not influenced by a nonspecific motor effect.

**Figure 3.**
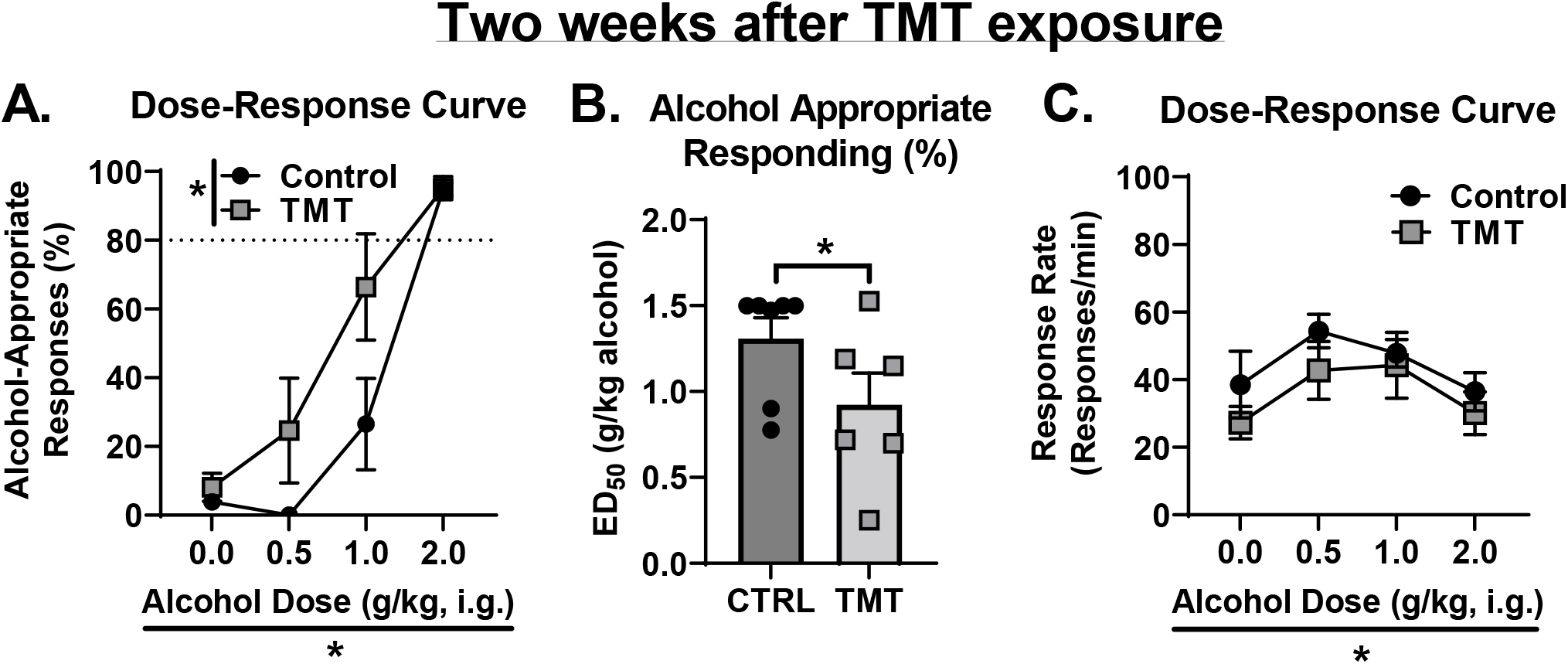
(A) 2 weeks after stressor, the TMT group showed increased alcohol-appropriate responses. (B) ED_50_ of the dose-response curve for alcohol-appropriate responses was lower in the TMT group. (C) TMT did not affect response rate, but alcohol did affect response rate. Dotted lines at 80% indicate full substitution for the 2 g/kg alcohol training dose. * p<0.05

#### Molecular adaptations of TMT exposure in the anterior insular cortex and the nucleus accumbens

The results for the c-Fos (Experiment 2) and gene expression (Experiment 3) experiments are shown in Fig. 4. The TMT group showed increased c-Fos immunoreactivity in the anterior insular cortex (aIC, Fig. 4A, t(13)=2.80, p=0.01) and the nucleus accumbens core (AcbC, Fig. 4B, t(13)=2.79, p=0.02) compared to controls 100 min after TMT exposure. These data show that TMT exposure results in neuronal activation in the aIC and AcbC.

**Figure 4.**
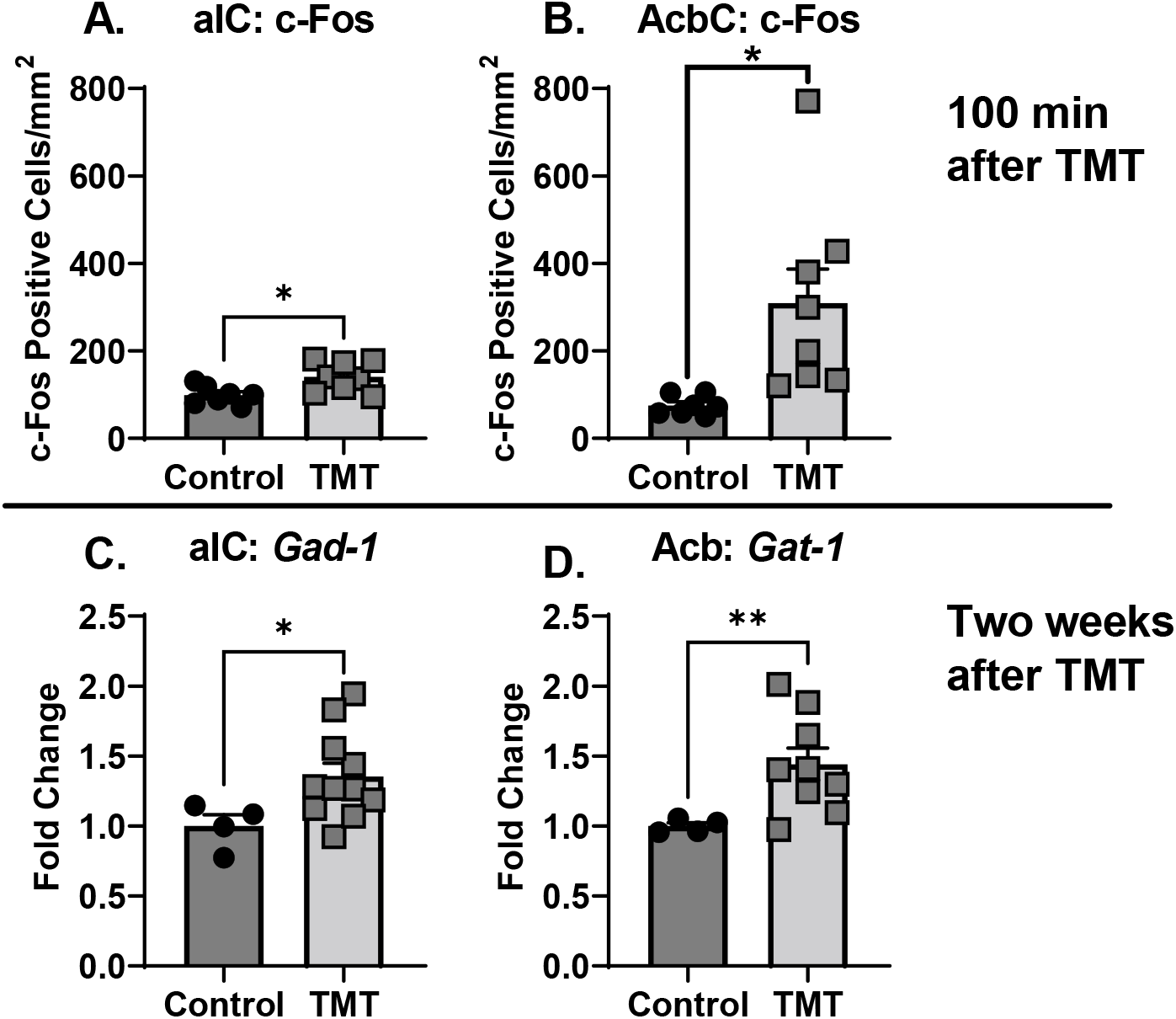
Molecular adaptations of TMT exposure in the anterior insular cortex and the nucleus accumbens. The TMT group showed increased c-Fos immunoreactivity compared to controls in the (A) aIC and (B) AcbC. The TMT group showed increased gene expression of *Gad-1* in the (C) aIC and *Gat-1* in the (D) Acb compared to controls. * p<0.05

Two weeks after TMT exposure (Experiment 3), in the anterior insular cortex (aIC) *Gad-1* gene expression was increased in the TMT group (Fig. 4C, t(10.76)=2.83, p=0.01). *Gad-2* expression did not differ between TMT and control group but was trending towards an increase in expression (p=0.07; CTRL: 1.0 ± 0.16; TMT: 1.48 ± 0.18). *Gat-1* was not tested in the aIC due to an unfortunate loss of samples. In the Acb (Fig. 4D), *Gat-1* was elevated as a result of TMT exposure (t(8.69)=3.73, p=0.005). *Gad-1* (CTRL: 1.0 ± 0.068; TMT: 1.5 ± 0.28) and *Gad-2* (CTRL: 1.0 ± 0.12; TMT: 1.11 ± 0.10) expression were not affected by TMT exposure in the Acb.

#### TMT exposure produces stress-reactive behaviors in alcohol-naïve and alcohol-experienced rat cohorts during the TMT exposure

Rats engaged in numerous stress-reactive behaviors during the TMT exposure. These behaviors during the TMT exposure for Experiments 1 and 3 are illustrated in Figure 5. In rats trained on the operant alcohol discrimination (Experiment 1 – alcohol-experienced), TMT exposure increased time spent digging (Fig. 5A, t(11)=5.11, p=0.0003), but did not affect time spent immobile (Fig. 5B). TMT exposure decreased time spent grooming (Fig. 5C, t(11)=3.60, p=0.004) and increased distance traveled (Fig. 5D, t(11)=5.83, p=0.0001). TMT exposure did not affect the time spent on the TMT side (Fig. 5E) but increased the number of midline crossings (Fig. 5F, t(11)=2.98, p=0.012) compared to controls.

**Figure 5.**
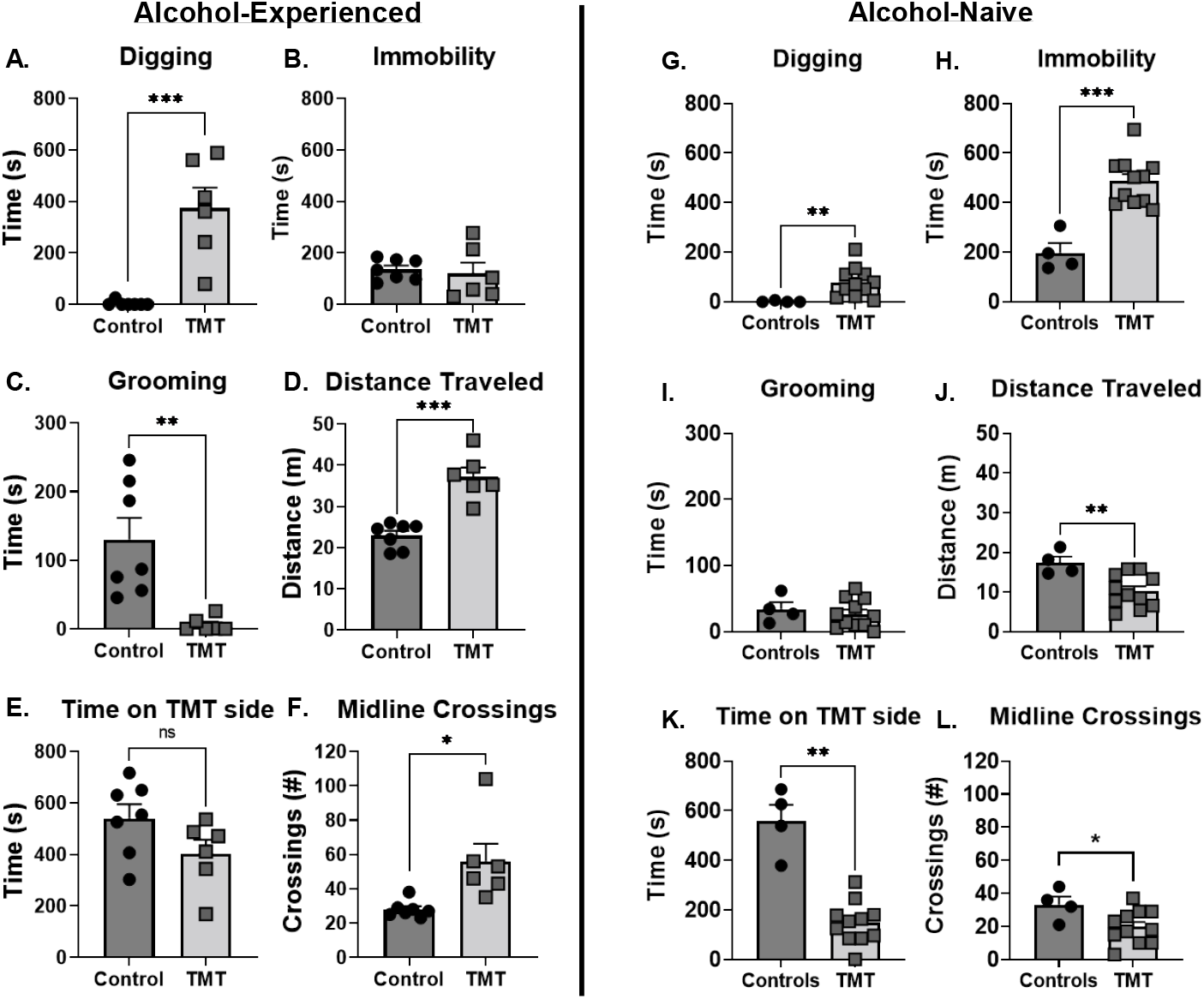
Stress-reactive behaviors in alcohol-experienced and alcohol-naïve cohorts during the TMT exposure. Stress reactive behaviors during the TMT exposure for the alcohol-experienced cohort (Experiment 1, A-F). (A) TMT exposure increased time spent digging, but (B) did not affect immobility. (C) TMT exposure decreased grooming behavior. (D) TMT increased distance traveled and (E) did not affect time spent on the TMT side. (F) TMT increased the number of midline crossings. Stress reactive behaviors during the TMT exposure for the alcohol-naive cohort (Experiment 3, G-L). (G) TMT exposure increased digging and (H) immobility. (I) TMT did not affect grooming. (J) TMT exposure decreased distance traveled, (K) time spent on the TMT side, and (L) the number of midline crossings.

In alcohol-naïve rats (Experiment 3, gene expression), TMT exposure increased time spent digging (Fig. 5G, t(10.18)=4.22, p=0.002) and time spent immobile (Fig. 5H, t(6.76)=5.97, p=0.0006). TMT exposure did not affect time spent grooming (Fig. 5I), but decreased distanced traveled (Fig. 5J, t(7.37)=3.69, p=0.007), time spent on the TMT side (Fig. 5K, t(3.92)=5.73, p=0.005) and the number of midline crossings (Fig. 5L, t(5.65)=2.38, p=0.05) compared to controls. While behavior from Experiment 1 and 3 were not statistically compared, the alcohol discrimination group appears to show a different behavioral response to TMT relative to the alcohol-naïve group.

## Discussion

These studies show that TMT exposure produces potentiated sensitivity to the interoceptive effects of alcohol 2 weeks after the exposure. Molecular experiments demonstrate neuronal activation as measured by c-Fos immunoreactivity immediately following TMT, and increased GABAergic gene expression 2 weeks after TMT in the anterior insular cortex (aIC) and nucleus accumbens (Acb). Together, these data indicate that TMT exposure results in lasting potentiation of alcohol interoceptive sensitivity, possibly through GABAergic adaptations in the aIC and Acb. Additionally, the degree of engagement in stress reactive behaviors during the TMT exposure differed in alcohol-naïve and alcohol-experienced rats, suggesting a potential influence of alcohol experience.

We initially hypothesized that TMT exposure would blunt interoceptive sensitivity to alcohol [43]. This hypothesis was supported by previous experiments showing blunted interoceptive sensitivity to alcohol following 7-day exposure to corticosterone in the rats’ drinking water [47–49]. However, contrary to this hypothesis, we found *potentiated* sensitivity to the interoceptive effects of alcohol 2 weeks after the TMT exposure. As such, these differences may be a consequence of the type of stressor-related models used (i.e., single exposure to an ethologically relevant stressor - predator odor as in the present study vs. a physiological stressor – corticosterone for 7 days). Note that these studies also differ in the alcohol training dose (1 g/kg vs. 2 g/kg) and pretreatment time of alcohol administration (10 min vs. 20 min) used, both of which may also play a role in the different effects observed. The present data are the first to suggest the possibility that different stress models may have opposite effects on alcohol interoceptive sensitivity.

The anterior insular cortex (aIC) and the nucleus accumbens core (AcbC) showed neuronal activation during TMT exposure as evidenced by increased c-Fos immunoreactivity. 2 weeks after the stressor, gene expression data showed increased expression of *Gad-1* in the aIC and *Gat-1* in the Acb. *Gad-1* is a marker of GABAergic cells, which suggests that TMT exposure increased GABAergic cell number in the aIC. *Gat-1* is the gene that encodes the GABA neurotransmitter transporter protein (GAD-1), which suggests altered GABA neurotransmitter levels in the synapse in the Acb. Prior data have shown increased NMDA receptor gene expression in the aIC 2 weeks after TMT [60], suggesting increased excitatory signaling in the aIC. As such, increased *Gad-1* expression in the aIC observed here may be the result of compensatory increase of local, GABAergic inhibition onto excitatory cells to restore changes in excitatory/inhibitory ratio [65]. GABA receptor potentiation by alcohol in the aIC and Acb both play a key role in the interoceptive effects of alcohol [14, 15, 53, 54]. As such, the GABAergic adaptations in the aIC and Acb 2 weeks after TMT exposure may in part underlie the potentiated sensitivity to alcohol.

We did not conduct statistical comparisons between the alcohol-experienced, discrimination-trained (Experiment 1) and alcohol-naïve (Experiment 3) rats because they are from separate cohorts, and there are several differences other than alcohol experience between groups (i.e., daily gavage, behavioral training, and food restriction in the alcohol-discrimination group). Nevertheless, visual comparison of groups shows some striking differences that may suggest that alcohol experience affects behavioral responses to a predator odor stressor. Both cohorts showed increased digging behavior in the TMT group, consistent with previous work [39, 60]. However, the alcohol-experienced cohort engaged in digging behavior to a greater extent than the alcohol-naïve cohort. Additionally, the alcohol-experienced cohort did not show increased immobility (which in part captures freezing, see [39]) during the TMT exposure, whereas the alcohol-naïve cohort did. Immobility during TMT exposure is consistent with prior work [39, 40]. Time spent grooming during the TMT exposure was decreased in alcohol-experienced rats but unchanged in alcohol-naïve rats. In relation to distanced traveled during the TMT exposure, the alcohol-experienced rats showed an increase, whereas the alcohol-naïve rats showed a decrease, likely because the naïve group was engaging in immobility behavior. The alcohol-experienced group did not show a difference in the time spent on the TMT side, whereas the alcohol-naïve rats spent significantly less time on the TMT side, likely because rats primarily engage in immobility behavior away from the TMT source (i.e., non-TMT side), which would lead to less time on the TMT side [39]. Finally, alcohol-experienced rats showed increased midline crossings during TMT exposure, whereas the alcohol-naïve group showed decreased midline crossings. Overall, the alcohol-experienced cohort (the discrimination cohort) showed a more hyperactive response during TMT (greater digging, decreased grooming, increased distance traveled and midline crossings, and no avoidance of the TMT side or immobility) than the alcohol-naïve rats. Alcohol-naive rats showed behavior more consistent with a fear response in rodents (i.e. freezing and avoidance) than the alcohol-experienced group. As such, alcohol experience may shift the behavioral response to a predator odor from that of freezing and avoidance to hyperactivity and increased active engagement (e.g., digging, increased distance traveled and midline crossings, decreased grooming). Another study using bobcat urine as the predator odor exposure showed that rats with a 5-week history of 2-bottle choice intermittent access to a 20% alcohol solution demonstrated a different stress response to the predator odor than an alcohol-naïve cohort. This model used classification criteria based on the time spent in proximity to the predator odor source during a context re-exposure test to sub-group rats into “avoiders” and “non-avoiders.” Specifically, alcohol experience was associated with a greater proportion of rats who met criteria for Avoiders [66], which is the sub-group that shows increased alcohol self-administration [44]. Together, these data show that a history of alcohol consumption prior to predator odor exposure may alter the behavioral consequences of the stressor. However, to make definitive conclusions based on direct comparisons between the groups, we would need to run a parallel cohort to the alcohol discrimination group that undergoes a similar behavioral training without alcohol.

Some limitations of these experiments should be addressed. First, these experiments were conducted in male rats only, and there may be important and clinically relevant sex differences in the consequences of predator odor exposure on sensitivity to the interoceptive effects of alcohol. Indeed, our lab has found differences in response to TMT exposure on alcohol-self-administration in males and females [40]. Secondly, gene expression and c-Fos experiments were conducted in alcohol-naïve rats, but the potentiated alcohol sensitivity effect was observed in an alcohol-experienced group, potentially affecting the way in which TMT exposure impacts molecular changes. It will be important for future work to examine these molecular changes in discrimination-trained rats. Finally, a functional connection between GABAergic adaptations and potentiated alcohol sensitivity was not made. Therefore, it will be interesting for future work to directly investigate this by testing substitution of a GABAergic positive modulator. Despite these limitations, these data are the first to show that predator odor exposure potentiates sensitivity to the interoceptive effects of alcohol.

These data demonstrate that TMT exposure has long-lasting (2 weeks) effects on alcohol interoceptive sensitivity, specifically potentiation of alcohol effects in male rats. Furthermore, we found increased expression of GABA-related genes in the aIC and Acb, which may underlie the potentiated alcohol sensitivity [7, 13–18, 51, 52]. Because the potentiation of the interoceptive effects of alcohol may increase alcohol consumption by priming continued drinking, these findings may have important implications for understanding PTSD-AUD comorbidity.

## Data Availability Statement

Data available on request from the authors.

